# Host-encoded CTCF regulates human cytomegalovirus latency via chromatin looping

**DOI:** 10.1101/2023.09.18.557586

**Authors:** Ian J. Groves, Christine M. O’Connor

**Affiliations:** Infection Biology Program, Global Center for Pathogen and Human Health Research, Lerner Research Institute, Cleveland Clinic, Cleveland, OH 44195; Molecular Medicine, Cleveland Clinic Lerner College of Medicine of Case Western Reserve University, Cleveland Clinic, Cleveland, OH 44195

**Keywords:** cytomegalovirus, CMV, latency, epigenetics, CTCF

## Abstract

Human cytomegalovirus (HCMV) is a prevalent pathogen that establishes life-long latent infection in hematopoietic cells. While this infection is usually asymptomatic, immune dysregulation leads to viral reactivation, which can cause significant morbidity and mortality. However, the mechanisms underpinning reactivation remain incompletely understood. The HCMV major immediate early promoter (MIEP)/enhancer is a key factor in this process, as its transactivation from a repressed to active state helps drive viral gene transcription necessary for reactivation from latency. Numerous host transcription factors bind the MIE locus and recruit repressive chromatin modifiers, thus impeding virus reactivation. One such factor is CCCTC-binding protein (CTCF), a highly conserved host zinc finger protein that mediates chromatin conformation and nuclear architecture. However, the mechanisms by which CTCF contributes to HCMV latency were previously unexplored. Here, we confirm CTCF binds two convergent sites within the MIE locus during latency in primary CD14^+^ monocytes, and following cellular differentiation, CTCF association is lost as the virus reactivates. While mutation of the MIE enhancer CTCF binding site does not impact viral lytic growth in fibroblasts, this mutant virus fails to maintain latency in myeloid cells. Furthermore, we show the two convergent CTCF binding sites allow looping to occur across the MIEP, supporting transcriptional repression during latency. Indeed, looping between the two sites diminishes during virus reactivation, concurrent with activation of MIE transcription. Taken together, our data reveal that three-dimensional chromatin looping aids in the regulation of HCMV latency, and provides insight into promoter/enhancer regulation that may prove broadly applicable across biological systems.

**Significance Statement:** Human cytomegalovirus (HCMV) remains an important healthcare consideration driven by disease in at-risk populations associated with reactivation of this virus from latent infection. We show here the establishment of latency is aided by a host nuclear architectural protein, CTCF. By binding two convergent sites on the virus major immediate early promoter/enhancer region, which largely acts as a switch from latency to reactivation, CTCF anchors a chromatin loop such that the virus promoter is maintained in a transcriptionally repressed state. Upon differentiation of cells, CTCF protein levels decrease, and this loop is alleviated as the virus reactivates. Our findings reveal further insight into the regulation of HCMV latency and promoter/enhancer elements, which is broadly applicable across biological systems.

## Introduction

Human cytomegalovirus (HCMV) remains an important healthcare issue world-wide. As the prototypic betaherpesvirus, HCMV infects between 44% to 96% of the world’s population, with local seroprevalence levels highly linked to socioeconomic variables (1, 2). As with all herpesviruses, HCMV infection is life-long, residing in a latent state in the host. While primary infection rarely causes disease in healthy individuals, reactivation from latency can lead to severe disease in immunosuppressed individuals, including transplant patients (3). Additionally, primary infection of the immunonaïve causes congenital infection, which can lead to symptoms ranging from benign to those that include hearing loss, developmental impairments, and microcephaly, to name a few (4). HCMV latency is maintained within cells of the myeloid lineage through the epigenetic control of viral transcription. During viral latent infection, a limited number of viral transcripts are synthesized, although major immediate early (MIE) gene expression is repressed (5). Transcription from the MIE locus is driven by the MIE promoter (MIEP), as well as additional alternative promoters that function in specific contexts (6), and activation of these promoters is regulated through chromatinization, as well as various transcription factors (7, 8). During reactivation, however, the MIEP becomes derepressed as the virus switches to a lytic infection and is able to fully replicate (5).

Myriad repressive and activatory host-derived transcription factors bind the MIE locus (7); nevertheless, our understanding of the transcriptional regulation of this region remains incomplete. Recently, Elder et al. showed expression of host CCCTC-binding protein (CTCF) is upregulated and binds the MIE enhancer during HCMV latency in CD14^+^ cells (9), consistent with MIE locus repression. CTCF is a highly conserved zinc finger protein that acts as a transcriptional and epigenetic insulator and mediates higher order chromatin conformation and looping in the nucleus (10). CTCF influences gene expression by many viruses, both during latent or persistent infections, as well as during reactivation and lytic infection (11). CTCF also binds numerous insulator regions on the herpes simplex virus (HSV) genome, and importantly, dissociation of CTCF from these sites is necessary for reactivation (12-14). Additionally, CTCF is enriched at sites across gammaherpesviruses genomes, including Epstein Barr virus (EBV), where it functions during latency and reactivation (reviewed in (15)). Mechanistically, CTCF aids in chromatin loop formation, differentially regulating transcription from certain genomic regions dependent on the stage of infection (15). Along with various other host factors (16), CTCF anchors loops through the interaction of two CTCF proteins that bind sites convergent to one another on the same DNA strand (10). Intriguingly, in addition to the CTCF binding site in the MIE enhancer (9), there is a second site in the MIE locus Intron A (17). During lytic infection, CTCF association at this site is consistent with inhibition of MIE-driven transcription and overall viral replication (17). Pertinently, these two CTCF binding sites face one another, meaning these sites are convergent. Hence, we hypothesized CTCF regulates the local three-dimensional genome organization during HCMV infection.

Herein we show CTCF binds both the MIE enhancer and Intron A sites during CD14^+^ cell latent infection. Upon M-CSF treatment, which differentiates the cells and allows for virus reactivation, CTCF protein levels decrease, concomitant with reduced binding to the MIE locus. Mutation of the MIE enhancer binding site abrogated CTCF binding, which resulted in a lytic-like infection, despite latent culture conditions. Moreover, we show CTCF binding to both the enhancer and Intron A sites is consistent with loop formation across the MIE locus during latency, which is lost upon cellular differentiation and viral reactivation. Taken together, our data reveal CTCF regulates MIE-driven transcription through the formation of a three-dimensional chromatin loop, which is dynamically reversed during reactivation.

## Results

### CTCF is enriched at the MIE locus during CMV latency

Earlier work revealed CTCF binds the enhancer (9) and the first intron within the MIE locus (17) (Fig. 1A). To confirm these findings in our latency model systems, we mock- or latently-infected THP-1 cells (Fig. 1B-D) or CD14^+^ monocytes (Fig. 1E-G) with wild type TB40/E-*mCherry* (WT) for 7 days (d), after which we treated a portion of each culture with differentiation stimuli for a further 2d. We then quantified *UL123* transcription (Fig. 1B and E) and IE1 protein expression (Fig. 1C and F), both of which increase following differentiation stimuli. In latently-infected cells, CTCF protein levels increase with WT infection, consistent with previous findings (9), which then decrease upon cellular differentiation (Fig. 1C and F). We also found significant enrichment of CTCF at both the Intron A and MIE enhancer sites (Fig. 1A) in both WT-infected THP-1 (Fig. 1D) and CD14^+^ (Fig. 1G) cells, although the efficiency of binding is enhanced at the enhancer site, irrespective of cell type (Figs. 1D and G). Additionally, following differentiation of the cells and derepression of the MIE locus, CTCF enrichment significantly decreases (Figs. 1D and G), suggesting CTCF is lost as the cells differentiate and the virus reactivates.

**Figure 1.**
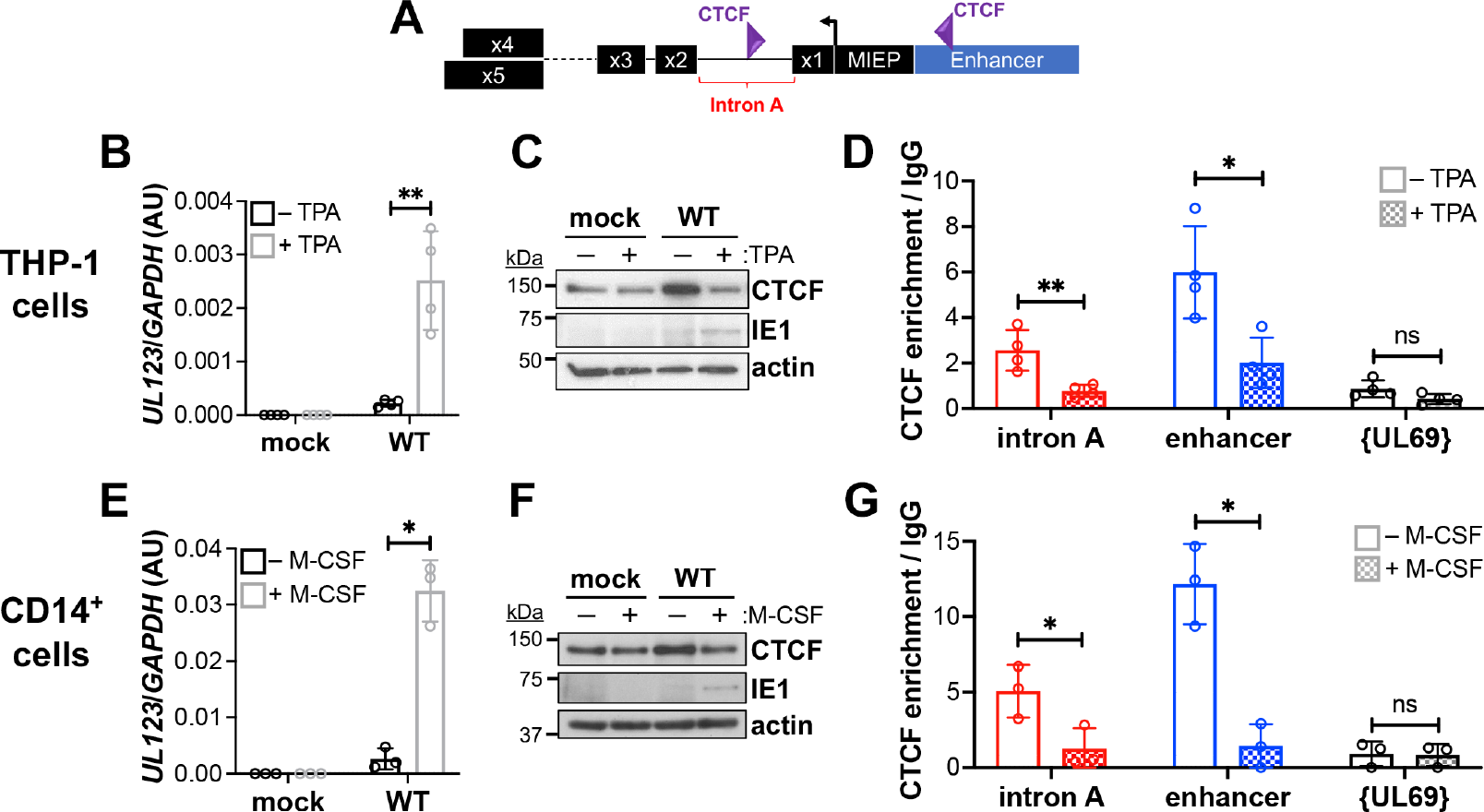
CTCF is enriched at Intron A and MIEP during HCMV latency in THP-1 and CD14^+^ monocytic cells. (**A**) Schematic of the CTCF binding sites in the MIE locus. CTCF is depicted as a gray triangle. Each exon, as well as Intron A, the MIEP, and the enhancer element are shown. (**B-D**) THP-1 cells or (**E-G**) CD14^+^ monocytes were mock- or WT-infected (MOI = 1.0 TCID_50_/cell) under latent conditions. At 7 dpi, cells were cultured an additional 2 d (**A-C**) with vehicle (DMSO; - TPA) or TPA (+TPA) or (**D-F**) under latent conditions (- M-CSF) or in the presence of the differentiation stimulus, M-CSF (+M-CSF). (**B, E**) *UL123* transcripts were quantified by RT-qPCR, and (**C, F**) host CTCF and CMV-encoded IE1 expression were assessed by immunoblot. Actin is shown as a loading control. Representative blots shown, n = 3. (**D, G**) CTCF enrichment was quantified at each MIE CTCF binding site. Data is shown as enrichment relative to the IgG control. The UL69 non-promoter region is shown as a negative control. (**B, D, E, G**) Data points (open circles) represent the mean of three technical replicates. Error bars indicate SD of the mean of three biological replicates. \**p*<0.05, \*\**p*<0.01, ns = not significant.

Since our data indicate CTCF is enriched at the MIE locus during latent infection yet lost upon differentiation of the cells as the virus reactivates, we hypothesized perhaps CTCF functions to maintain MIE repression during latency. To address this, we next designed a mutation at the enhancer binding site to prevent CTCF association (*SI Appendix*, Fig. S1A and B). We compared the MIE enhancer CTCF binding site to the predicted metazoan consensus sequence (18) and found it was in fact longer than previously recognized (9) (*SI Appendix*, Fig. S1B). Additionally, our computational analyses of this region using PhysBinder (19) revealed a putative CREB binding site within the CTCF binding site sequence (*SI Appendix*, Fig. S1B). Since phosphorylated CREB binding to the MIE locus is critical for reactivation (20), we designed our mutation such that this putative CREB binding site is retained (*SI Appendix*, Fig. S1B). To this end, we mutated nucleotides only within the highest region of homology with the consensus sequence to produce TB40/E-*mCherry*-CTCF*mut* (CTCF*mut*; *mut*), using previously described BAC recombineering technology (21). We then reverted the mutated nucleotides to the wild type sequence, generating TB40/E-*mCherry*-CTCF*rev* (CTCF*rev*; *rev*). Following sequence validation of each newly-generated virus, we then evaluated their lytic growth in NuFF-1 fibroblasts by performing a multi-step growth curve. CTCF*mut* displayed wild type growth kinetics, as we observed no significant difference in either cell-free (*SI Appendix*, Fig. S1C) or cell-associated virus (*SI Appendix*, Fig. S1D) compared to WT or CTCF*rev*, suggesting the mutation does not impact HCMV lytic replication.

### CTCF enrichment at the MIE locus is critical for maintaining viral latency

We next evaluated the contribution of CTCF binding to the enhancer site to viral latency. First, to confirm the enhancer binding site mutation disrupts CTCF association, we assessed CTCF enrichment using chromatin immunoprecipitation (ChIP). To this end, we infected either THP-1 or CD14+ cells with WT, CTCF*mut*, or CTCF*rev* under latent conditions for 7 d, after which a portion of each infected culture was treated with differentiation stimuli for an additional 2 d. CTCF was indeed enriched at the enhancer binding site in cells latently-infected with WT or CTCF*rev*, which was lost upon cellular differentiation (Fig. 2A and E), similar to our above observations (Fig. 1D and G). In contrast, the CTCF*mut*-infected cells cultured under latent conditions failed to enrich CTCF at the enhancer site (Fig. 2A and E), without impacting the required recruitment of phosphorylated CREB (20) to the locus (*SI Appendix*, Fig. S1B) during reactivation (Fig. 2B and F). Since we hypothesize CTCF binding during latency impacts the activity of the MIE locus, we evaluated both *UL123* transcription and IE1 protein expression. As expected, WT and CTCF*rev* cells infected under latent conditions display low levels of *UL123* mRNA, while CTCF*mut* infected cells cultured under the same conditions had significantly higher expression levels of *UL123* transcript (Fig. 2C and G). Consistent with these data, IE1 is robustly expressed in CTCF*mut* cells infected under latent conditions, which increases in all infected populations following differentiation of these cells (Fig. 2D and H), suggesting CTCF*mut*-infected cells favor a lytic-like infection. Supporting these findings, *UL123* (Fig. 2C and G) and IE1 (Fig. 2D and H) expression in the CTCF*mut*-infected cultures maintained under latent conditions are similar to levels observed following differentiation of the WT- or CTCF*rev*-infected cells. Collectively, these data suggest CTCF binding to the MIE enhancer site is critical for maintaining HCMV latency.

**Figure 2.**
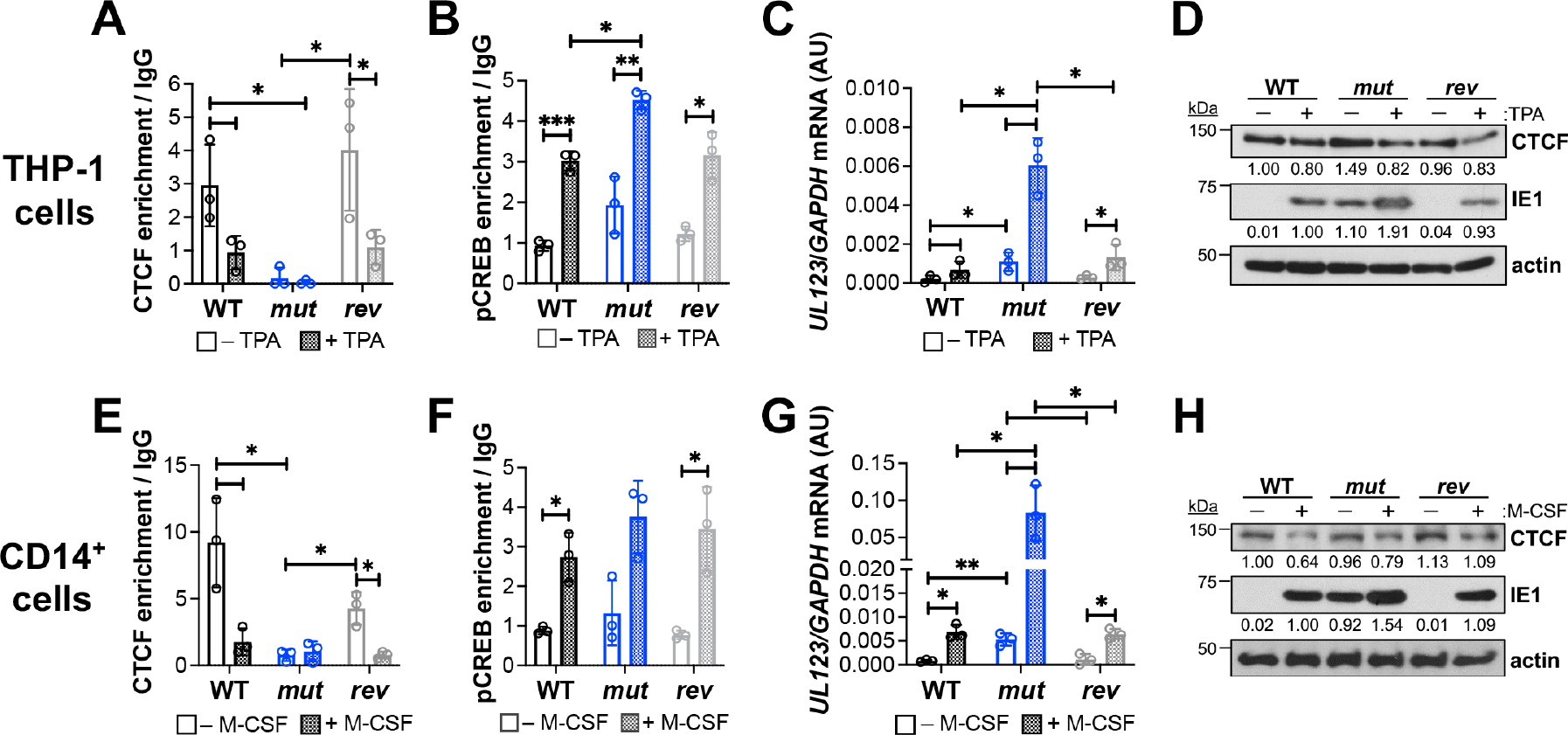
Mutation of the CTCF binding site in the MIE enhancer results in derepression of the MIE locus. (**A-D**) THP-1 or (**E-H**) CD14^+^ cells were infected (MOI = 1.0 TCID_50_/cell) under latent conditions with WT, CTCF*mut* (*mut*), or CTCF*rev* (*rev*). At 7 dpi, cells were cultured an additional 2 d (**A-D**) with vehicle (DMSO; -TPA) or TPA (+TPA), or (**E-H**) under latent conditions (- M-CSF) or in the presence of the differentiation stimulus, M-CSF (+M-CSF). (**A-B, E-F**) Enrichment of (**A, E**) CTCF or (**B, F**) phosphorylated CREB (pCREB) at the MIE enhancer binding site was assessed by ChIP and is depicted as enrichment relative to the IgG control. (**C, G**) *UL123* mRNA levels were quantified by RT-qPCR and are plotted relative to cellular *GAPDH*. (**D, H**) Cellular CTCF, HCMV IE1, and actin (loading control) expression were evaluated by immunoblot. Representative blots shown, n = 3. (**A-C, E-G**) Data points (open circles) represent the mean of three technical replicates. Error bars indicate SD of the mean of three biological replicates. \**p*< 0.05, \*\**p*<0.01, \*\*\**p*<0.001.

Our data suggest CTCF binding to its site within the MIE enhancer is important for repression of the MIE locus. Thus, to determine if this impacts the maintenance of viral latency, we infected CD14^+^ monocytes with WT, CTCF*mut*, or CTCF*rev* under latent conditions for 7 d, after which a portion of each infected culture was maintained under these conditions or treated with M-CSF to differentiate the cells, thus reactivating the virus. We then determined the frequency of infectious centers by extreme limiting dilution analysis (ELDA; (22)). During latency, WT- and CTCF*rev*-infected cells display a low frequency of infectious centers, which significantly increase following reactivation (Fig. 3 and *SI Appendix* Fig. S2), as expected. In contrast, CTCF*mut*-infected cells display a significant increase in the frequency of infectious centers relative to that of WT- or CTCF*rev*-infected cultures, despite their maintenance in medium favoring latency (Fig. 3, open blue bars). Indeed, the CTCF*mut*-infected CD14^+^ cells treated with M-CSF exhibit a further significant increase in the frequency of infectious centers (Fig. 3 and *SI Appendix* Fig. S2, checkered blue bars), which is more than 8-fold higher than control viruses in differentiated cells (Fig. 3 and *SI Appendix* Fig. S2, checkered black or gray bars), likely due to the fact CTCF*mut*-infected cells already favored a more lytic-like infection at the time of M-CSF addition. In sum, these data confirm CTCF binding to the MIE enhancer site is required for efficient viral latency.

**Figure 3.**
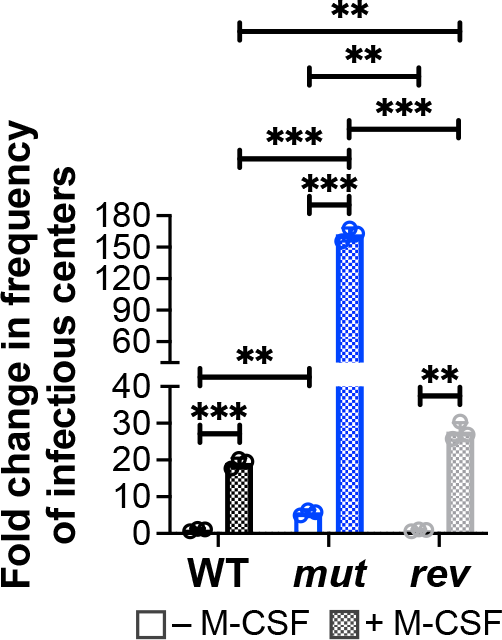
Mutation of the CTCF binding site in the MIE enhancer results in increased virus replication in CD14^+^ monocytes. CD14^+^ monocytes were infected with WT, CTCF*mut* (*mut*), or CTCF*rev* (*rev*) (MOI = 1.0 TCID_50_/cell) under latent conditions. At 7 dpi, cells were either maintained under latent conditions (- M-CSF) or treated with the differentiation stimulus, M-CSF (+ M-CSF) and co-cultured with naïve NuFF-1 cells to quantify the frequency of infectious centers by ELDA. Each data point (circle) is the mean of three technical replicates (see *SI Appendix*, Fig. S2). Data is shown as fold change relative to WT latent (- M-CSF) culture conditions (open black bar). Error bars indicate SD of three biological replicates. \**p*<0.05, \*\**p*<0.01, \*\*\**p*<0.001.

### CTCF mediates looping between convergent binding sites in the MIE locus

Thus far, our findings reveal CTCF association with the MIE enhancer binding site impacts MIEP activity and viral latency. Further, our data show CTCF is also enriched at the Intron A site, though to lesser affinity than the enhancer site (Fig. 1D and G). Importantly, the Intron A site faces the enhancer site (Fig. 1A), consistent with a structure that has the potential to function in looping of the DNA (23). Furthermore, CTCF controls transcription from other virus genomes (24), thus there is precedence for CTCF-mediated regulation of HCMV gene expression. Taken together, this led us to hypothesize that looping, anchored by CTCF bound at the Intron A and MIE enhancer binding sites during latency may, at least in part, control transcription from this locus. To test this, we employed chromatin conformation capture (3C) (25) to determine if these two loci are closer in three-dimensional (3D) space than would otherwise be expected from a linear DNA strand. Briefly, THP-1 or CD14^+^ cells were infected with WT, CTCF*mut*, or CTCF*rev* (MOI = 1.0 TCID_50_/cell) for 7 d. We then either maintained the infected cells under latent conditions or treated them with differentiation stimuli for an additional 2 d, after which we assessed the 3D architecture of the genome by 3C-PCR (Fig. 4A). As controls, we retained a portion of the fixed cells prior to digesting the DNA (input), as well as following the enzymatic digestion step (digest). We then amplified the resulting ligated product, if any, by PCR. As expected, using PCR primers that span the Intron A and enhancer CTCF binding sites resulted in the amplification of a 1009 bp product in all ‘input’ samples, whereas there was no product in our ‘digested’ samples. However, in our ‘ligated’ samples, PCR amplification resulted in 136 bp products in the WT- and CTCF*rev*-latently-infected cells (Fig. 4B and F). The presence of this band was reduced in CTCF*mut*-infected cells cultured under latent conditions and was diminished further to undetectable levels following differentiation of either infected cell type. This is in comparison to low remaining levels of this 136 bp product in either WT- and CTCF*rev*-infected cells treated with differentiation stimuli. While these data suggest the genome is looping in WT, latently-infected cells, we were unable to detect the cohesion subunit, structural maintenance of chromosome protein 3 (SMC3), at the MIE locus by ChIP (*SI Appendix*, Fig. S3), which is often, but not always, necessary for maintenance of CTCF-associated genomic loops (26). To ensure DNA was loaded equally into each reaction, we amplified a *NlaIII*-insensitive region (150 bp) of the MIE locus (Fig. 4C and G). Finally, to confirm specificity of the 3C-PCR primers for the region spanning the two CTCF binding sites, we Sanger sequenced products from the WT-infected THP-1 cell ‘input’ and ‘ligated’ samples, thus confirming the DNA junction within the ‘input’ fraction (Fig. 4D), as well as a shorter, chimeric sequence in the ligated sample (Fig. 4E), consistent with re-ligation of two usually separated sequences. Together, these data indicate CTCF mediates a loop between its Intron A and enhancer binding sites.

**Figure 4.**
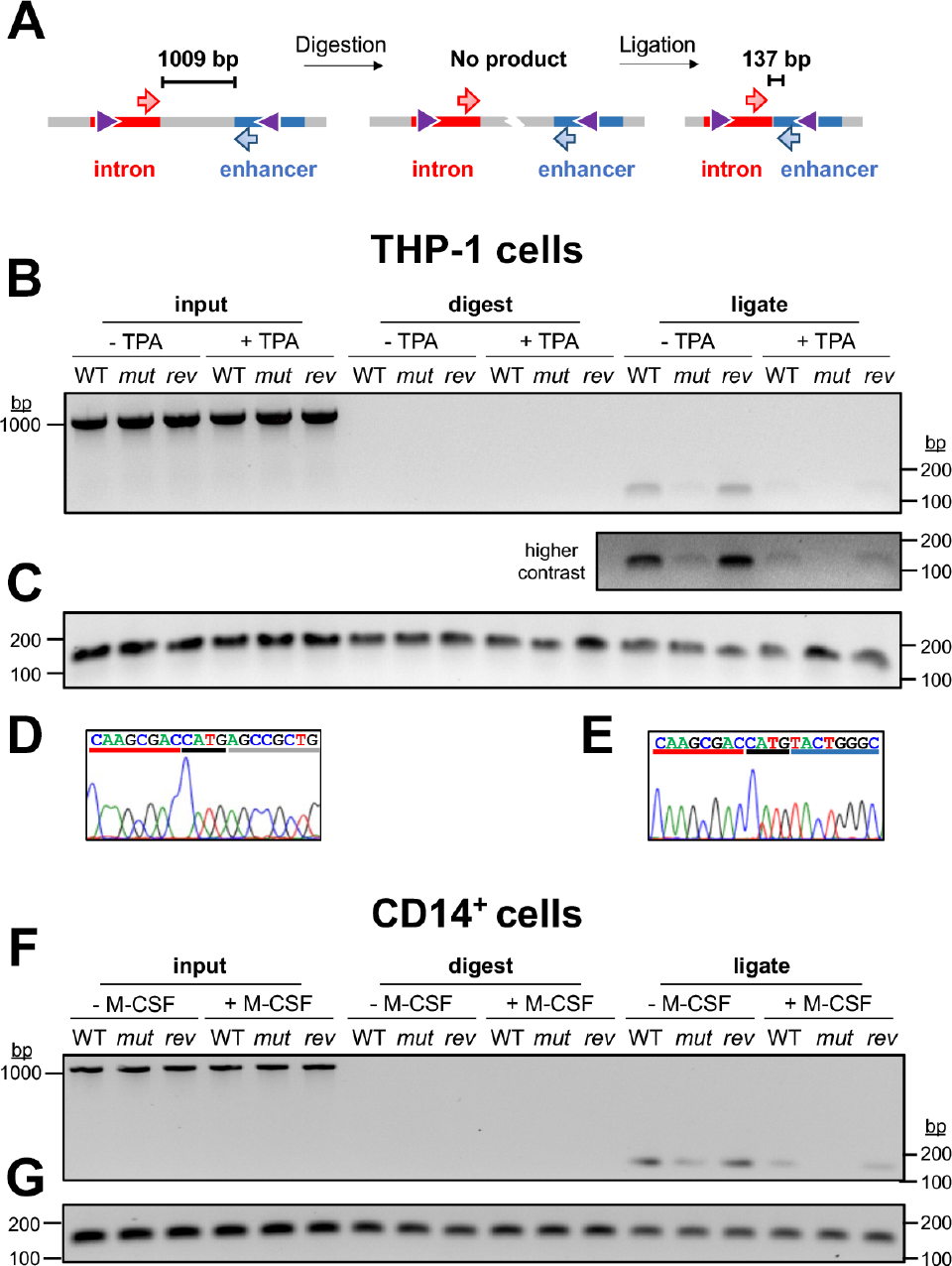
Abrogation of CTCF binding at the enhancer binding site decreases chromatin looping. (**A**) Schematic of chromatin conformation capture (3C) –PCR technique. Purple triangles, CTCF. PCR primers, orange/green arrows. (**B-E**) THP-1 or (**F, G**) CD14^+^ cells were infected (MOI = 1.0 TCID_50_/cell) with WT, CTCF*mut* (*mut*), or CTCF*rev* (*rev*) under latent conditions. At 7 dpi, cells were cultured an additional 2 d (**B-E**) with vehicle (DMSO; -TPA) or TPA (+TPA), or (**F, G**) under latent conditions (-M-CSF) or in the presence of the differentiation stimulus, M-CSF (+M-CSF). Cells were then fixed with formaldehyde and chromatin conformation capture (3C) was performed, as described in detail in the Materials and Methods. (**B, F**) PCR analysis of 3C products using primers spanning the possible looping loci: a 1009 bp product (uncut template, experimental loading control), 136 bp product (ligation of digested ends without the intermediary sequence). (**C, G**) Amplification of a 150 bp region within a sequence that is not cut with *Nla*III is shown as a PCR loading control. (**D, E**) PCR products from (**B**) were excised from gels, purified, and Sanger sequenced to show (**D**) a wild type sequence and (**E**) a chimeric sequence. *Nla*III site is underlined in black; sequence in the intronic and enhancer regions are also underlined in red and blue, respectively. (**B, C, F, G**) Representative gels shown, n = 3.

## Discussion

Herein we show CTCF binding at the MIE locus during HCMV latency is critical for repression of this region. Specifically, our data reveal CTCF binds convergent sites in the enhancer and Intron A (Fig. 1). Further, mutation of the enhancer binding site, which limits CTCF association at the enhancer, results in a lytic-like infection in monocytic cells cultured under latent conditions (Fig. 2), and consistent with this, CTCF*mut*-infected cells fail to maintain viral latency (Fig. 3). Finally, our data reveal CTCF mediates repression of the MIE locus by mediating looping between its two convergent sites in the MIE locus (Fig. 4). Collectively, our findings reveal another layer by which the MIE region is regulated during latency and reactivation.

Convergent CTCF binding sites act as boundaries in the metazoan genome, leading to looping of chromatin across megabase-wide regions, thereby resulting in topologically-associating domains (TADs). Additionally, CTCF-mediated looping occurs more locally between promoters and enhancers (10). Viral gene expression modulated via CTCF looping also occurs and plays important function in the lifecycle of many double-stranded DNA viruses, including alpha- and gamma-herpesviruses (11). Our findings herein now add to our understanding of the function of CTCF during herpesvirus infection and represent the first report of CTCF-mediated control of betaherpesvirus regulation. While our data indicate CTCF binding to the enhancer site is critical for maintaining latency (Fig. 3), mutation of the Intron A CTCF binding site showed a 10-fold increase in viral replication during lytic infection of permissive cells (17), indicating the intronic site acts to repress transcription from this locus, possibly by affecting RNA polymerase II function. Although the impact of the Intron A CTCF binding site in models of HCMV latency remain untested, it is attractive to hypothesize mutation of this site would similarly lead to the inability of the convergent sites to loop the DNA, resulting in derepression of the MIE locus.

What other host and or viral factors might influence CTCF-mediated repression of the MIE locus? Based on previous work (9) and our findings herein, we propose a model whereby US28-stimulated upregulation of CTCF protein expression (9) results in CTCF enrichment at, and looping of, the MIE locus in latently-infected myeloid cells. Chromatin loops are often anchored by CTCF through extrusion of the DNA/protein complex through a cohesin structure, until the point at which the convergent CTCF proteins interact (10). While we show the association of CTCF with the MIE locus is consistent with chromatin looping (Fig. 4), we were unable to detect cohesin binding in this region (*SI Appendix*, Fig. S3). Perhaps this is not surprising; cohesin-CTCF complexes selectively anchor long-range loops, whereas loops formed by sequences in close range are likely anchored by CTCF alone (27). Whether another cellular or viral factor function in cohesin’s place is currently unknown. Nevertheless, the loss of this loop during reactivation of the virus is consistent with concomitant differentiation-dependent decreases in CTCF levels. Indeed, CTCF association with the two MIE binding sites is likely aided by a US28-mediated increase in CTCF protein levels (28). During latency, US28 signaling is also important for attenuating the function of transcriptional activators, such as AP-1, to support maintenance of MIE-driven transcriptional repression (29, 30). Moreover, assessing genome-wide chromatin interactions during differentiation of THP-1 cells to macrophages using Hi-C highlighted that AP-1 is integral for directing post-differentiation AP1-bound activation hubs at key macrophage genes (31), indicating chromatin loops are both stabilized or diminished via transcription factor modification. Further, HCMV-encoded GPCR UL33 signaling mediates CREB activation, which ameliorates MIE repression and facilitates, at least in part, viral reactivation (20). Intriguingly, one of these cAMP response elements (CREs, 5’-TGACGTCA-3’) to which CREB binds resides within the extended CTCF binding sequence in the MIEP enhancer (*SI Appendix*, Fig. S1B). Hence, it is possible that CREB activation, in addition to attenuated CTCF protein levels, tips the balance from the binding of repressive factors to those that are activators of the MIE locus, resulting in complete viral reactivation.

Taken together, our findings show the association of host CTCF protein at the MIE locus during HCMV latency, at least in part, functions to repress MIE-driven transcription by forming a repressive chromatin loop between the enhancer and Intron A CTCF anchor sites. Loss of this loop during differentiation of cells is concurrent with reduced binding of CTCF and derepression of the MIE locus, consistent with reactivation of the virus. Whether this form of gene expression control is found further across the HCMV genome during latency/reactivation and/or lytic infection requires further studies, as those recently performed for gammaherpesviruses EBV (32) and KSHV (33). Moreover, whether additional host and/or viral proteins bind or interact with CTCF at these sites (16) play a part during latency, as shown for 3D control of transcription of other viruses (34-36), remains unanswered. In addition to AP-1, BRD4 also affects chromatin looping during differentiation (37). Understanding the involvement of BRD4 in this mechanism is critical, as this bromodomain protein represents a potential therapeutic target for HCMV infection (38). Thus, further inroads into our understanding of how virus gene expression is controlled through mechanisms such as these could improve future HCMV-specific therapies.

## Methods

### Cells and Viruses

Methods for cell culture, propagation of viruses, and generation of viral recombinants are included in *SI Appendix, SI Methods*.

### Latency and Reactivation Assays

Experimental details for these assays are in *SI Appendix, SI Methods*.

### RNA and Protein Analyses

Details regarding analyses of RNA and protein are included in *SI Appendix, SI Methods*.

### Multistep Growth Analyses

Experimental approaches for analyzing viral growth are in *SI Appendix, SI Methods*.

### Extreme Limiting Dilution Assay (ELDA)

Reactivation efficiency of HCMV infection in CD14^+^ monocytes was measured by ELDA, essentially as described previously (22). Briefly, CD14^+^ cells were latently infected for 7 d and then either maintained under latent conditions or cultured in reactivation medium. Cultures were then serially diluted two-fold onto naïve NuFF-1 cells and cultured for an additional 14 d. The production of infectious virus was quantified using viral-expressed mCherry as a marker of infection and ELDA software (bioinf.wehi.edu.au/software/elda/index.html).

### Chromatin Immunoprecipitation (ChIP)

THP-1 cells infected as indicated above were then fixed in 1.0% formaldehyde for 15 min at RT, prior to quenching with 125 mM glycine for 5 min. Cells were lysed in ChIP buffer (150 mM NaCl; 50 mM Tris-HCl, pH 7.5; 5 mM EDTA; 0.5% Igepal-CA630; 1.0% Triton X-100; protease inhibitors [Roche]) and debris was removed by centrifugation. Chromatin was then sheared to 0.3-to 1.0-kb fragments using a Diagenode Bioruptor Pico (30 s on/30 s off, 12 cycles (38) and aliquots stored as input controls. CTCF was immunoprecipitated using protein A agarose (MilliporeSigma) and 2 µg of anti-CTCF antibody (Abcam, ab188408) or rabbit IgG isotype (Abcam, ab171870) as a negative control. DNA was captured by incubation with Chelex beads (BioRad), eluted by boiling, treated with proteinase K treatment (0.2 ng/μl), and finally purified using chromatography columns (BioRad). For CD14^+^ monocytes, sonicated samples were chromatin-immunoprecipitated using the ChIP-IT PBMC kits (Active Motif), according to the manufacturer’s instructions (38). The same sonication conditions and quantities of antibodies used for THP-1 cells were also used for processing the CD14^+^ cells. Enrichment of CTCF-bound DNA in both cell types was then quantified against rabbit isotype control (set to 1.0) using primers directed at the MIE locus or the UL69 non-promoter region as a negative control (*SI Appendix*, Table S2).

### Chromatin Conformation Capture (3C)-PCR

3C was performed essentially as previously described (34). Briefly, THP-1 or CD14^+^ cells (3x10^6^) were infected as described above and then washed in 1 x PBS and resuspended in 1.0 ml 10% FBS)/PBS. Cells were fixed in 10 ml total of 1.0% formaldehyde in 10% FBS/PBS for 10 min at RT before glycine (125 mM final concentration) treatment for 5 min. Cells were then pelleted (500 x *g*) at 4°C for 10 m, after which they were resuspended in 1.0 ml of ice-cold lysis buffer (10 mM Tris-HCl, pH 7.7; 10 mM NaCl; 5 mM MgCl2; 0.1 mM EGTA; protease inhibitors [Roche]) and incubated on ice for 10 min before nuclei were pelleted (400 × *g*) for 5 min at 4°C. Next, cell nuclei were resuspended in 500 μl of 1.2X restriction enzyme buffer and 0.3% SDS (final concentration) and incubated at 37°C with shaking at 300 rpm (Eppendorf Thermomixer). After 1 h, 50 μl of 20% Triton X-100 was added and samples were incubated at 37°C for an additional 1 h with shaking (300 rpm). At this point, 10% of each sample was retained as an input control, while 200 U of *NlaIII* were added to each remaining sample and incubated overnight at 37°C with shaking (300 rpm). Samples were then halved and 50% was retained as a digest control. SDS (0.1% final concentration) was added to the remaining half of each sample and then incubated for 25 min at 65°C with shaking (300 rpm). Next, 6.5 ml of 1.15X ligation buffer containing Triton X-100 (1% final concentration) was added, and samples were incubated for 1 h at 37°C with gentle shaking (100 rpm). T4 DNA ligase (100 U) was then added, and samples were incubated for 4 h at 16°C, followed by 30 min at RT. All input controls, digest controls, and ligation samples were then treated with 30 µg proteinase K for 16 h at 65°C, followed by 300 µg RNase A for 30 min at 37°C. DNA was then isolated by phenol:chloroform extraction and alcohol precipitation, and all samples were resuspended in 10 mM Tris-HCl, pH 7.5 for use in PCR. All 3C primers are listed in *SI Appendix*, Table S2.

## Notes

### Competing Interest Statement

The authors have declared no competing interest.

## References

1. M. Zuhair et al., Estimation of the worldwide seroprevalence of cytomegalovirus: A systematic review and meta-analysis. Rev Med Virol 29, e2034 (2019).

2. K. Fowler et al., A systematic literature review of the global seroprevalence of cytomegalovirus: possible implications for treatment, screening, and vaccine development. BMC Public Health 22, 1659 (2022).

3. P. Griffiths, M. Reeves, Pathogenesis of human cytomegalovirus in the immunocompromised host. Nat Rev Microbiol 19, 759–773 (2021).

4. R. F. Pass, R. Arav-Boger, Maternal and fetal cytomegalovirus infection: diagnosis, management, and prevention. F1000Res 7, 255 (2018).

5. S. M. Matthews, I. J. Groves, C. M. O’Connor, Chromatin control of human cytomegalovirus infection. mBio 10.1128/mbio.00326-23, e0032623 (2023).

6. D. Collins-McMillen, J. Kamil, N. Moorman, F. Goodrum, Control of Immediate Early Gene Expression for Human Cytomegalovirus Reactivation. Front Cell Infect Microbiol 10, 476 (2020).

7. A. L. Dooley, C. M. O’Connor, Regulation of the MIE Locus During HCMV Latency and Reactivation. Pathogens 9 (2020).

8. D. Collins-McMillen, J. Buehler, M. Peppenelli, F. Goodrum, Molecular Determinants and the Regulation of Human Cytomegalovirus Latency and Reactivation. Viruses 10 (2018).

9. E. G. Elder, B. A. Krishna, E. Poole, M. Perera, J. Sinclair, Regulation of host and viral promoters during human cytomegalovirus latency via US28 and CTCF. J Gen Virol 102 (2021).

10. B. Dehingia, M. Milewska, M. Janowski, A. Pekowska, CTCF shapes chromatin structure and gene expression in health and disease. EMBO Rep 23, e55146 (2022).

11. C. S. Varghese, J. L. Parish, J. Ferguson, Lying low-chromatin insulation in persistent DNA virus infection. Curr Opin Virol 55, 101257 (2022).

12. M. K. Ertel, A. L. Cammarata, R. J. Hron, D. M. Neumann, CTCF occupation of the herpes simplex virus 1 genome is disrupted at early times postreactivation in a transcription-dependent manner. J Virol 86, 12741–12759 (2012).

13. S. D. Washington et al., Depletion of the Insulator Protein CTCF Results in Herpes Simplex Virus 1 Reactivation In Vivo. J Virol 92 (2018).

14. J. S. Lee et al., CCCTC-Binding Factor Acts as a Heterochromatin Barrier on Herpes Simplex Viral Latent Chromatin and Contributes to Poised Latent Infection. mBio 9 (2018).

15. L. B. Caruso, D. Maestri, I. Tempera, Three-Dimensional Chromatin Structure of the EBV Genome: A Crucial Factor in Viral Infection. Viruses 15 (2023).

16. X. Sun, J. Zhang, C. Cao, CTCF and Its Partners: Shaper of 3D Genome during Development. Genes (Basel) 13 (2022).

17. F. P. Martinez et al., CTCF binding to the first intron of the major immediate early (MIE) gene of human cytomegalovirus (HCMV) negatively regulates MIE gene expression and HCMV replication. J Virol 88, 7389–7401 (2014).

18. A. V. Persikov, M. Singh, De novo prediction of DNA-binding specificities for Cys2His2 zinc finger proteins. Nucleic Acids Res 42, 97–108 (2014).

19. S. Broos et al., PhysBinder: Improving the prediction of transcription factor binding sites by flexible inclusion of biophysical properties. Nucleic Acids Res 41, W531–534 (2013).

20. B. A. Krishna, A. B. Wass, A. L. Dooley, C. M. O’Connor, CMV-encoded GPCR pUL33 activates CREB and facilitates its recruitment to the MIE locus for efficient viral reactivation. J Cell Sci 134 (2021).

21. C. M. O’Connor, T. Shenk, Human cytomegalovirus pUS27 G protein-coupled receptor homologue is required for efficient spread by the extracellular route but not for direct cell-to-cell spread. J Virol 85, 3700–3707 (2011).

22. M. Peppenelli, J. Buehler, F. Goodrum, Human Hematopoietic Long-Term Culture (hLTC) for Human Cytomegalovirus Latency and Reactivation. Methods Mol Biol 2244, 83–101 (2021).

23. E. de Wit et al., CTCF Binding Polarity Determines Chromatin Looping. Mol Cell 60, 676–684 (2015).

24. I. Pentland, J. L. Parish, Targeting CTCF to Control Virus Gene Expression: A Common Theme amongst Diverse DNA Viruses. Viruses 7, 3574–3585 (2015).

25. J. Dekker, K. Rippe, M. Dekker, N. Kleckner, Capturing chromosome conformation. Science 295, 1306–1311 (2002).

26. J. Zuin et al., Cohesin and CTCF differentially affect chromatin architecture and gene expression in human cells. Proc Natl Acad Sci U S A 111, 996–1001 (2014).

27. S. Sofueva et al., Cohesin-mediated interactions organize chromosomal domain architecture. EMBO J 32, 3119–3129 (2013).

28. M. S. Humby, C. M. O’Connor, Human Cytomegalovirus US28 Is Important for Latent Infection of Hematopoietic Progenitor Cells. J Virol 90, 2959–2970 (2015).

29. B. A. Krishna, A. B. Wass, C. M. O’Connor, Activator protein-1 transactivation of the major immediate early locus is a determinant of cytomegalovirus reactivation from latency. Proc Natl Acad Sci U S A 117, 20860–20867 (2020).

30. B. A. Krishna, M. S. Humby, W. E. Miller, C. M. O’Connor, Human cytomegalovirus G protein-coupled receptor US28 promotes latency by attenuating c-fos. Proc Natl Acad Sci U S A 116, 1755–1764 (2019).

31. D. H. Phanstiel et al., Static and Dynamic DNA Loops form AP-1-Bound Activation Hubs during Macrophage Development. Mol Cell 67, 1037–1048 e1036 (2017).

32. S. M. Morgan et al., The three-dimensional structure of Epstein-Barr virus genome varies by latency type and is regulated by PARP1 enzymatic activity. Nat Commun 13, 187 (2022).

33. M. Campbell et al., KSHV Topologically Associating Domains in Latent and Reactivated Viral Chromatin. J Virol 96, e0056522 (2022).

34. I. Pentland et al., Disruption of CTCF-YY1-dependent looping of the human papillomavirus genome activates differentiation-induced viral oncogene transcription. PLoS Biol 16, e2005752 (2018).

35. D. Li, T. Mosbruger, D. Verma, S. Swaminathan, Complex Interactions between Cohesin and CTCF in Regulation of Kaposi’s Sarcoma-Associated Herpesvirus Lytic Transcription. J Virol 94 (2020).

36. L. N. Lupey-Green et al., PARP1 Stabilizes CTCF Binding and Chromatin Structure To Maintain Epstein-Barr Virus Latency Type. J Virol 92 (2018).

37. R. Linares-Saldana et al., BRD4 orchestrates genome folding to promote neural crest differentiation. Nat Genet 53, 1480–1492 (2021).

38. I. J. Groves et al., Bromodomain proteins regulate human cytomegalovirus latency and reactivation allowing epigenetic therapeutic intervention. Proc Natl Acad Sci U S A 118 (2021).

