## Supplemental Files for "Host-encoded CTCF regulates human cytomegalovirus latency via chromatin looping"

### Supporting Information Text

#### SI Methods

**Cells and Viruses**. Primary human newborn foreskin fibroblasts (NuFF-1 cells, GlobalStem [passages 13 – 29]) were maintained in Dulbecco's modified Eagle medium (DMEM), supplemented with 10% fetal bovine serum (FBS), 2 mM L-glutamine, 0.1 mM non-essential amino acids, 10 mM HEPES, and 100 U/ml of penicillin and streptomycin. THP-1 cells (ATCC) were cultured in Roswell Park Memorial Institute (RPMI)-1640 medium supplemented with 10% FBS and 100 U/ml of penicillin and streptomycin and maintained at densities between $3\times10^5$ and $8\times10^5$ cells/ml. Primary $CD14^+$ monocytes were isolated from de-identified cord blood samples (Abraham J. & Phyllis Katz Cord Blood Foundation d.b.a. Cleveland Cord Blood Center & Volunteer Donating Communities in Cleveland and Atlanta) by magnetic separation, as described elsewhere (1) and were cultured minimally at $3x10^6$/ml in low serum medium (XVIVO-15; Lonza), supplemented with 2 mM L-glutamine. All cells were maintained at 37°C and 5% CO2.

Bacterial artificial chromosome (BAC) recombineering using the clinical CMV strain TB40/E (clone 4), previously engineered to express mCherry to monitor infection (TB40/E*mCherry*, referred to herein as WT) (2), was used as a template to generate TB40/E*mCherry*-CTCF*mut* using galactokinase (galK) recombineering, as described elsewhere (3). Essentially, the galK gene was PCR amplified from pGalK by using primers listed in SI Appendix, Table S1. Recombination-competent SW105 Escherichia coli containing TB40/E*mCherry* were electroporated with the resulting product. The mutation in the MIE enhancer CTCF binding site was introduced using a gBlock, which was amplified by PCR (Table 1). GalK-positive clones were transformed using this product, after which clonse were counter-selected against galK. The resulting virus mutant, TB40/E*mCherry*-CTCF*mut* (CTCF*mut*) was then used as a template to repair the mutated residues to the wild type sequence, ultimately creating TB40/E*mCherry*-CTCF*rev* (CTCF*rev*). The genomic accuracy of viruses was verified by Sanger sequencing.

All viral stocks were propagated on and titered by 50% tissue culture infectious dose ($TCID_{50}$) on naïve NuFF-1 fibroblasts, as described elsewhere (2).

**Latency and Reactivation Assays.** THP-1 cells, pre-cultured in X-VIVO-15 ($5\times10^5$/ml) for 48 hours (h), were infected at a multiplicity of infection (MOI) of 1.0 TCID50/cell by centrifugal enhancement (1000 x *g* for 30 minutes (min) at room temperature (RT)) in X-VIVO-15 and then incubated for 60 min at 37°C and 5% $CO_2$. Viral inocula were removed, cells washed three times with 1× PBS, and then re-plated in X-VIVO-15. After 7 days (d), cells were treated with 12-O-tetradecanoylphorbol-13-acetate (TPA) to induce cellular differentiation and viral reactivation or vehicle control (DMSO) for an additional 2 d.

CD14+ monocytes ($3x10^6$/ml) were similarly infected by centrifugal enhancement (1000 x *g*, 30 min, RT), then incubated at 37°C/5% $CO_2$ for 16 h. Cells were then washed three times in 1× PBS and cultured in X-VIVO-15. At 7 d, a portion of each infected cell population was cultured in reactivation media (RPMI-1640, containing 10% FBS, with macrophage colony-stimulating factor (M-CSF, 10 ng/ml)) or maintained under conditions favoring latency (X-VIVO-15) for a period of time dependent on the individual experiment.

**RNA and Protein Analyses.** Total RNA was isolated from cells using the High Pure RNA Isolation Kit (Roche), as per the manufacturer's instructions. RNA (1.0 µg per sample) was then reverse transcribed (RT) using the TaqMan RT reagent kit (Applied Biosystems) with random hexamers. Viral *UL123* transcript level was assessed by quantitative polymerase chain reaction (qPCR) using SYBR green PCR mix (Applied Biosystems) and quantified relative to host *GAPDH* transcript levels. Primers for RT-qPCR analyses are shown in SI Appendix, Table S2.

For protein analyses, cells were lysed in radioimmunoprecipitation assay (RIPA) buffer (1.0% NP-40; 1.0% sodium deoxycholate; 0.1% SDS; 0.15 M NaCl; 0.01 M $NaPO_4$, pH 7.2; 2.0 mM EDTA) on ice for 1 hour, vortexing every 15 min. Total protein concentrations were quantified by Bradford assay using Protein Assay Dye Reagent Concentrate (Bio-Rad). Samples were then denatured at 95°C for 10 min before equal quantities of protein per sample were separated by SDS–polyacrylamide gel electrophoresis (SDS-PAGE) and transferred to 0.45 µm nitrocellulose (Amersham) by semi-dry transfer. Proteins were detected by immunoblot using the following antibodies: anti-CMV IE1 (clone 1B12, 1:50)(4), anti–β-actin peroxidase (MilliporeSigma, 1:20,000), and goat anti-mouse, goat anti-rat, and goat anti-rabbit HRP secondary antibodies (all from Jackson ImmunoResearch Labs; 1:10,000). Relative protein levels were then quantified by densitometry using ImageJ (5).

**Multistep Growth Analyses.** Assessment of virus replication was performed by infecting NuFF-1 cells at a MOI of 0.01 $TCID_{50}$/cell. Cell-free viral supernatants were collected over a 16 d time course, at which point cell-associated virus was also harvested. All samples were stored at −80°C until processing. Infectious virus was then titrated on naïve NuFF-1 cells and analyzed by $TCID_{50}$ assay.

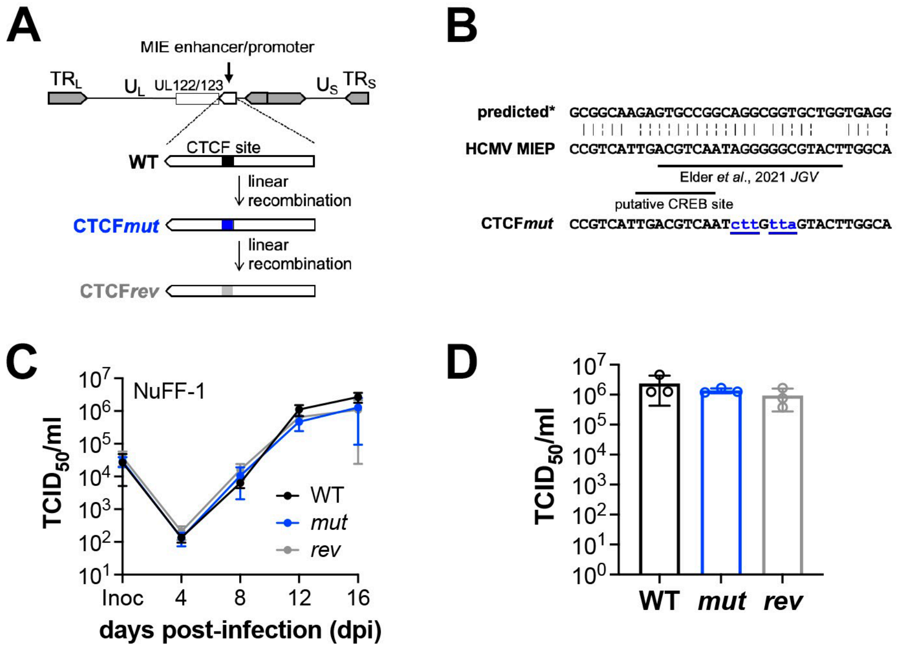


**Fig. S1. CTCF*mut* grows with wild type kinetics in lytically-infected fibroblasts.** (**A**) BAC-derived TB40/E-*mCherry* (WT) was used to generate a mutation in the CTCF binding site (black box) of the MIE enhancer/promoter region, termed TB40/E-*mCherry*-CTCF*mut* (CTCF*mut*, blue box). The mutated nucleotides (lowercase letters underlined in blue in (**B**) were then repaired to the wild type sequence, yielding the revertant virus, TB40/E-*mCherry*-CTCF*rev* (CTCF*rev*). (**B**) The MIEP CTCF binding site sequence for WT HCMV is depicted aligned with a previously published metazoan predicted consensus sequence (ref. (6)); '|', nucleotide homology; '¦', base homology). Altered bases in CTCF*mut* are shown as lowercase letters underlined in blue. (**C**) Viruses in (**A**) were used to infect NuFF-1 fibroblasts to determine multistep growth (MOI = 0.01 $TCID_{50}$/cell) over a 16-day time course. Cell-free virus was collected at the times indicated, and (**D**) cell-associated virus was harvested at 16 dpi. (**C, D**) Viral titers were quantified by $TCID_{50}$ using naïve NuFF-1 fibroblasts. Samples were analyzed in triplicate and data are mean±SD of the biological replicates. No statistically significant difference between viruses was found at individual time points.

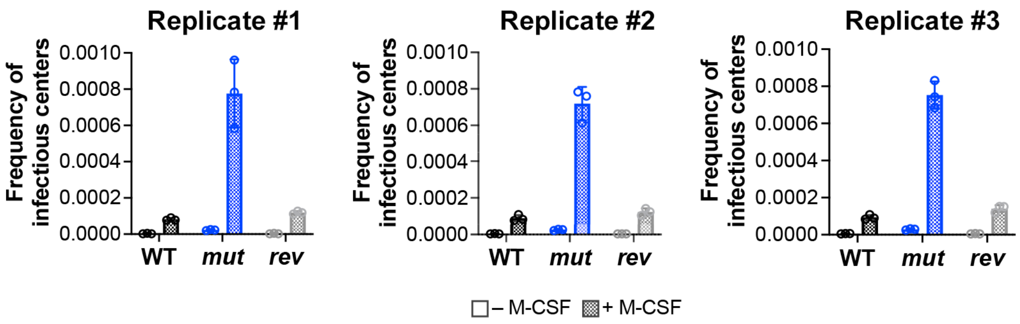


**Fig. S2. Biological replicates for the ELDA data presented in Figure 3.** CD14$^+$ cells were infected (MOI = 1.0 $TCID_{50}$/cell) with WT, CTCF*mut* (*mut*), or CTCF*rev* (*rev*) under latent conditions. At 7 dpi, cells were treated with vehicle (- M-CSF) or + M-CSF and co-cultured with naïve NuFF-1 cells to quantify the frequency of infectious centers by ELDA. Each data point (circles) represents an individual technical replicate within the experiment.

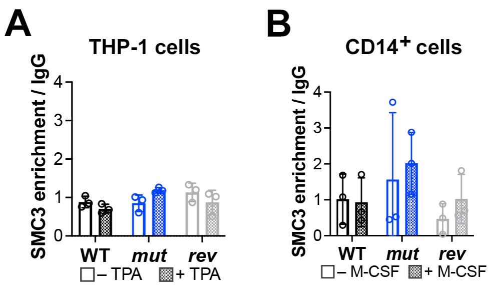


**Fig S3. SMC3, a Cohesin complex subunit, is not recruited to the MIE locus during infection.** (**A**) THP-1 or (**B**) $CD14^+$ cells were infected (MOI = 1.0 $TCID_{50}$/cell) under latent conditions with WT, CTCF*mut* (*mut*), or CTCF*rev* (*rev*). At 7 dpi, cells were cultured for an additional 2 d (**A**) with vehicle (DMSO; -TPA) or TPA (+TPA), or (**B**) under latent conditions (- M-CSF) or in the presence of the differentiation stimulus, M-CSF (+ M-CSF). (**A-B**) Enrichment of SMC3 at the MIE enhancer binding site was assessed by ChIP and is depicted as enrichment relative to the IgG control. Each data point (open circle) represents the mean of three technical replicates, and the error bars indicate the SD of the mean of three biological replicates.

**Table S1. Oligonucleotides used for generation of viral recombinants.**

| Primer Use | Sequence* | Target |
|---|---|---|
| *galK* insertion (CTCF*mut*-galK intermediate) | GAAAGTCCCGTAAGGTCATGTACTGGGCATAATGCCAGGCGGGCCATTTACCTGTTGACAATTAATCATCGGCA | MIEP CTCF–galK Fwd |
| | GGAGTATTTACGGTAAACTGCCCACTTGGCAGTACATCAAGTGTATCATATCAGCACTGTCCTGCTCCTT | MIEP CTCF–galK Rev |
| CTCF*mut* insert | AGGAAAGTCCCGTAAGGTCATGTACTGGGCATAATGCCAGGCGGGCCATTTACCGTCATTGACGTCAATcttGttaGTACTTGGCATATGATACACTTGATGTACTGCCAAGTGGGCAGTTTACCGTAAATACTCC | CTCF*mut* gBlock |
| CTCF*mut* insert and WT amplification (CTCF*mut* & CTCF*rev* generation) | AGGAAAGTCCCGTAAGGTCA | CTCF*mut* gBlock Fwd & WT |
| | TTACGGTAAACTGCCCACTTG | MIEP CTCF*mut* gBlock Rev & WT |
| Sequencing primers for WT, CTCF*mut*, CTCF*rev* | GACATTTTGGAAAGTCCCGTTG | MIEP seq Fwd |
| | TCCGCGTTACATAACTTACGGT | MIEP seq Rev |

*Primer sequences are shown 5' to 3' in orientation. Underlined sequences denote those corresponding to the pGalK plasmid. Lower case letters are mutation sites in the MIEP CTCF binding site.

**Table S2. Oligonucleotide pairs used for PCR.**

| Primer Use | Forward (5' to 3') | Reverse (5' to 3') | Target |
|---|---|---|---|
| RT-qPCR | ATTTTCTGGGCATAAGCCATAATC | GCCTTCCCTAAGACCACCAAT | *UL123* cDNA |
| RT-qPCR | CTGTTGCTGTAGCCAAATTCGT | ACCCACTCCTCCACCTTTGAC | *GAPDH* cDNA |
| ChIP-qPCR | CTGCTCAGACTACACTGCCC | TTAAGGCAGCGGCAGAAGAA | enhancer CTCF binding site |
| ChIP-qPCR | CGTAAGGTCATGTACTGGGCA | GCCCACTTGGCAGTACATCA | intronA CTCF binding site |
| ChIP-qPCR | GGCTGATGATCTTGCGGGAA | CGAGAGTCTACGTCTGGCAC | UL69 (control locus) |
| 3C-PCR | CTGGCCTCCACTGTTAGGAG | CCCACTTGGCAGTACATCAA | intronA-MIE region |
| 3C-PCR | AACAGCGTGGATGGCGTCTCC | GGCACCAAAATCAACGGGACTTT | MIE *NlaIII*-resistant locus |

**SI References**


1. E. Poole, I. Groves, S. Jackson, M. Wills, J. Sinclair, Using Primary Human Cells to Analyze Human Cytomegalovirus Biology. *Methods Mol Biol* **2244**, 51-81 (2021).
2. C. M. O'Connor, T. Shenk, Human cytomegalovirus pUS27 G protein-coupled receptor homologue is required for efficient spread by the extracellular route but not for direct cell-to-cell spread. *J Virol* **85**, 3700-3707 (2011).
3. S. Warming, N. Costantino, D. L. Court, N. A. Jenkins, N. G. Copeland, Simple and highly efficient BAC recombineering using galK selection. *Nucleic Acids Res* **33**, e36 (2005).
4. H. Zhu, Y. Shen, T. Shenk, Human cytomegalovirus IE1 and IE2 proteins block apoptosis. *J Virol* **69**, 7960-7970 (1995).
5. C. A. Schneider, W. S. Rasband, K. W. Eliceiri, NIH Image to ImageJ: 25 years of image analysis. *Nat Methods* **9**, 671-675 (2012).
6. A. V. Persikov, M. Singh, De novo prediction of DNA-binding specificities for Cys2His2 zinc finger proteins. *Nucleic Acids Res* **42**, 97-108 (2014).
